# Effects of Maleic Hydrazide on Germination, Radicle Growth and Mitotic Division of *Trigonella foenum-graecum* L. and *Allium cepa* L

**DOI:** 10.1101/2025.06.28.662147

**Authors:** A. A. El-Ghamery, Mohamed. A. Mousa

## Abstract

The effects of different treatments with maleic hydrazide which is an herbicide (growth retardant) on the cytology and growth of *Trigonella foenum-graecum* (fenugreek) and *Allium cepa* (onion) were investigated. Six concentrations of maleic hydrazide ranging from 5 to 55 ppm were applied for 6, 12, 18, 24, 36 and 48h. The treatments reduced the germination percentages of *Trigonella foenum-graecum* and *Allium cepa* and inhibited the root growth of both plants. Concentrations higher 50 ppm for 48h were toxic for both plants. Analysis of Variance (ANOVA) showed that there was significant difference (P < 0.05 and P < 0.01) in the mean root length of both plants exposed to different concentrations of the maleic hydrazide. This indicated that the root growth inhibition was concentration dependent.

The non-lethal of MH showed an inhibitory effect on cell division in root tips of both plants and caused a decrease in their mitotic index values. The reduction in MI in root tips of *Allium cepa* was more evident than that of *Trigonella foenum-graecum*. All treatments changed the frequency of mitotic phases as compared with the control values. All the used concentrations of MH significantly induced a number of chromosomal aberrations in root tip cells of *Trigonella foenum-graecum and Allium cepa*. The total percentages of abnormalities in *Allium cepa* root tip cells were more than that in *Trigonella foenum-graecum* with all concentrations less than 45 ppm of MH. The most dominant types of observed abnormalities were stickiness, C-mitosis and distributed metaphase and anaphase. MH treatments produced a number of mitotic abnormalities in dividing cells in root tips of both plants resulting from its action on the spindle apparatus such as C-metaphases, lagging chromosomes and multipolar anaphases and telophases. Also, MH induced vacuolated nuclei and irregular prophases. The induction of chromosomal stickiness and chromosomal aberrations such as bridges indicates its action on the chromosome. This abnormality (chromosomal bridges at Ana-telophases) indicates true clastogenic potential of the chemical. It may be concluded that MH causes toxic effect on root tip cells of *Trigonella foenum-graecum* and *Allium cepa* and this toxicity induces different types of genic and chromosomal variations.

## Introduction

It was indicated that many cytogenetic studies have been carried out to detect the harmful effect of different pesticides on different plants **(Marcano *et al*., 2004)**. These chemicals used at excessive dosages can give rise to abnormal chromosomes and degeneration in mitotic cycle, such as micronucleus formation, chromosome bridges, and polyploidy **(Tosun *et al*., 2001)** and in meiotic cell division **(Çali, 2009)**. Growth retardants have been proven to prevent excessive stem elongation and reduce internode length in plants by inhibiting the effect of cell division and enlargement of cell in plants **(Ganesh *et al*., 2014)**.

In early works the inhibitory effects of MH on plant growth were mainly considered to result from the suppression of plant metabolism (inhibition of enzyme activity) and interference of the compound with plant hormones and growth regulators. Numerous experiments performed with various plant species have shown that MH acts as an inhibitor of the synthesis of nucleic acids and proteins. Among higher plants, selective sensitivity to the toxic effects of MH is well proved. This phenomenon seems to be due to the differential ability of various plant species to detoxicate the chemical. Plants can break down MH into several products, one of which, hydrazine, is a well-known mutagen and carcinogen. The compound is degraded by soil micro-flora and hence can be utilized as a source of nitrogen nutrition **(Swietlińska and Žuk, 1978)**. Recently, the results have been obtained by **Pandy (2008)** showed that some of the agricultural chemicals are mitotic depressive in higher concentrations and mitotic promoter in lower concentrations and induced a variety of chromosomal abnormalities.

Maleic hydrazide (MH), chemically defined as 1,2-dihydro-3,6-pyridazinedione, is a structural isomer of uracil and is used in agriculture as a commercial herbicide to retard the development of buds so that vegetables last longer in storage and arrive fresher to the consumer **(Lee *et al*., 2001)**. MH is a clastogenic and mutagenic agent that may cause spindle fiber defects **(Rank *et al*., 2002)**, or inhibition of nucleic acids and proteins synthesis, and also regulation of auxin metabolism **(Ito *et al*., <u>2001</u>)**. When maleic hydrazide (MH) was applied an inhibition of uracil uptake was observed **(Coupland and Peel, 1972)**. Maleic hydrazide is claimed to be selectively toxic to plants but not to bacteria and fungi **(Epstein *et al*., 1967)**. Maleic hydrazide has long been as an inhibitor of plant growth, **Wiszniowski *et al*. (2009)** reported that the occurrence of growth inhibition of *Vicia faba* roots exposed to maleic hydrazide, which is non-hormonic herbicide used to inhibition plant growth a depressant of plant growth, suppression of grass growth and growth of shrubs and trees, and to stop sprouting of vegetables in storage. Moreover, it is used in agriculture in despite its known effect as a mutagenic and clastogenic agent **(Swietlińska and Žuk, 1978)**. Use in excess of some plant growth regulators and fertilizers in order to obtain higher yields of crops can cause genotoxic effects on living organisms **(Marcano *et al*., 2004)**. The effects of maleic hydrazide (MH) on various plants had been investigated such as *Rumex acetosa* (Mendhulkar, 2000), *Vicia faba* **(Patra *et al*., 2000; Miadokova *et al*., 2001; De Marco *et al*., 2005; Sobita and Bhagirath, 2005, Wiszniowski *et al*., 2009 and Laskar and Khan, 2014)**, *Trigonella foenum-graecum* **(Jabee *et al*., 2008, Siddiqui *et al*., 2012 and Hasan *et al*., 2018), Oryza *sativa* (Mendhulkar, 2000 and Das *et al*., 2008)**, *Triticum aestivum* **(Mendhulkar, 2000)**, *Allium cepa* **(Ferrara *et al*., 2001, Patra *et al*., 2005, Ghosh *et al*., 2010, Sabale and Mane, 2011)**, *Helianthus annuus* **(Kaymak, 2005)**, *Hordeum vulgare* **(Stroyev, 1968, Juchimiuk *et al*., 2007 and Kwasniewska *et al*., 2018)**, *Pisum sativum* **(Edwin and Reddy, 1993)**, *Picea abies* **(Schubert and Rieger, 1994)**, *Crepis capillaris* **(Juchimiuk and Maluszynska, 2005)**, *Nicotiana tabacum* **(Gichner *et al*., 2000 and Juchimiuk *et al*., 2006)**, *Albizia lebbeck* **(Tomar, 2008)**, *Cassia tora* **(Khosla and Dnyansagar, 1980)** and *Phelipanche aegyptiaca* **(Venezian *et al*., 2017)**.

The extensive use of herbicides in agriculture and their potential carcinogenicity strongly suggest the need to extend the cytotoxic evaluation of these compounds by using different methods. Therefore, the objective of the present study is to compare the different responses of *Allium cepa* L. (studies made by **Fiskesjo (1997)** pointed out that the *Allium* test is useful for the detection of potentially cyto-genotoxic substance) and *Trigonella foenum-graecum* L. to the effects of maleic hydrazide (MH), under the same condition, during seed germination, root elongation and mitotic activity then evaluation the mutagenicity and/or genotoxicity of this chemical in the light of induction of mitotic abnormalities and as well as discussing the mechanism of its action. Its mode of action in plants is not clear, although several hypotheses have been proposed and investigated, such as inhibition of cell division by mitotic disruption **(Ganesh *et al*., 2014)**. Others have suggested that MH acts as an anti-auxin, anti-gibberellin or regulator of auxin metabolism and other plant growth regulators **(Hoffman and Parups, 1964)**.

## Materials and Methods

### Materials

Experimental plant materials selected for the present investigation were (*Allium cepa* L.) and fenugreek (*Trigonella foenum-graecum* L.) and the chemical “Maleic hydrazide” (MH) was used to induce cytogenetic variability in the selected plant materials. Maleic hydrazide (MH): C_4_H_4_N_2_O_2_ is an alkylating agent and a carcinogen. Healthy and uniform size seeds of *Allium cepa* L. (onion) and *Trigonella foenum-graecum* L. (fenugreek) obtained from Field Crops Institute, Agriculture Research Centre, Giza, Egypt. Maleic hydrazide (MH) supplied from sigma. MH is dissolved in redistilled water and the applied concentrations were 5, 15, 25, 35, 45, and 55 ppm after a preliminary test which showed that concentrations higher than 55 ppm applied for 48h exerted toxic effect on cells.

## Methods

### Seed germination and radicle length

Seeds of both plants immersed in 5% NaOCl (Sodium hypochloride solution) for 5 min to avoid fungal invasion. The seeds of both plants presoaked in distilled water for 2h, then placed directly in different concentrations of MH, ranging 5 to 55 ppm for 6 treatment times (6-48h). For each treatment, a triplicate of 25 seeds was used. The treated seeds (25 for each treatment) were washed carefully with distilled water then transferred to Petri-dishes containing filter paper moistened with distilled water and allowed to germinate at room temperature 25 ± l°C for 3 days. Control seeds were treated with distilled water. Seeds are considered to be germinated when radicle has emerged approximately ≥ 2 mm. After recording germination counts, the percentage of seed germination was calculated on the basis of total number of seeds sown **(Scott *et al*., 1984): Germination (%)** = No. of seeds germinated / No. of seeds sown ×100

For radicle length, thirty of the germinated seeds were immersed in suitable amount of each tested concentrations of 5-55 ppm of MH for the different times of 6-48 hours. Similarly, 30 seedling roots were soaked in distilled water for the same period was run with each treatment as the control. Following the treatments, the treated seedlings together with each control samples were kept in the dark at 23-25°C, in order to minimize the fluctuation in the rate of cell division **(Evans *et al*., 1957)**. At the end of each treatment time after 5 days, the length of the radicle was measured. The relative reduction of radicle length was calculated as the percentage of the deviation from the control (T/C%).

### Cytological study

Ten germinated seeds, with radicle 2-3 cm long, were treated with different concentrations for different times. Control germinated seeds were placed in distilled water. After each treatment, the roots were cut off and immediately fixed in glacial acetic acid: absolute ethyl alcohol (1:3 v/v) for overnight, then hydrolyzed in 1 N HCl at 58–60°C for 10 min **(Qian, 2004)**. The root tips were stained by using Feulgen’s squash technique **(Darlington and La Cour, 1976)**. At least three slides for each treatment were examined to determine the mitotic index (MI) which was calculated as the percentage of dividing cells to the total number of cells examined. The frequency of each mitotic phase was calculated as the percentage of dividing cells in that stage to the total number of dividing cells examined. The same slides were analyzed for the percentage and types of chromosomal abnormalities in cells at each mitotic phase as well as non-dividing cells.

### Statistical analysis

The significance of differences between treatments and control on both mitotic index and the frequency of chromosomal abnormalities was evaluated statistically. Each treatment was made in three replicates. For statistical analysis, one-way ANOVA (Sigma Plot 13.0 software) SPSS was used to determine significance at p < 0.05 and p < 0.01 (Duncan, 1955).

## Results

The seed germination percentages in both tested plants presented in Table l show that the percentages were reduced by all applied concentrations as compared with the control values. The reduction in the germination percentages increased with increase of MH concentrations and treatment durations. The treatment with 45 ppm of MH for 36h was the most effective in reducing the control value of 90 ± 3.3% for seeds of *Trigonella foenum-graecum* to the value of 3.3±3.3%. But the treatment with 55 ppm of MH for 24h was the most effective in reducing the control value of 90 ±3.3% for seeds of *Allium cepa* to the value of 3.3± 3.3%. For both tested plants, the treatment with 55 ppm of MH for 36 and 48h completely inhibited germination of seeds. Also, the results in Table 1 show that the effects of MH on root growth of both test plants varied with both concentrations applied and treatment times. The inhibition of root growth for each treatment time increased with increasing MH concentration from 5 to 55 ppm. After 6 h treatment, the roots were more or less normal throughout the whole range of MH concentrations from 5 to 55 ppm, while the treatment for 36 and 48 h with 55 ppm was the most effective in retardation of root growth, which was statistically significant (p < 0.01).

**Table 1.**
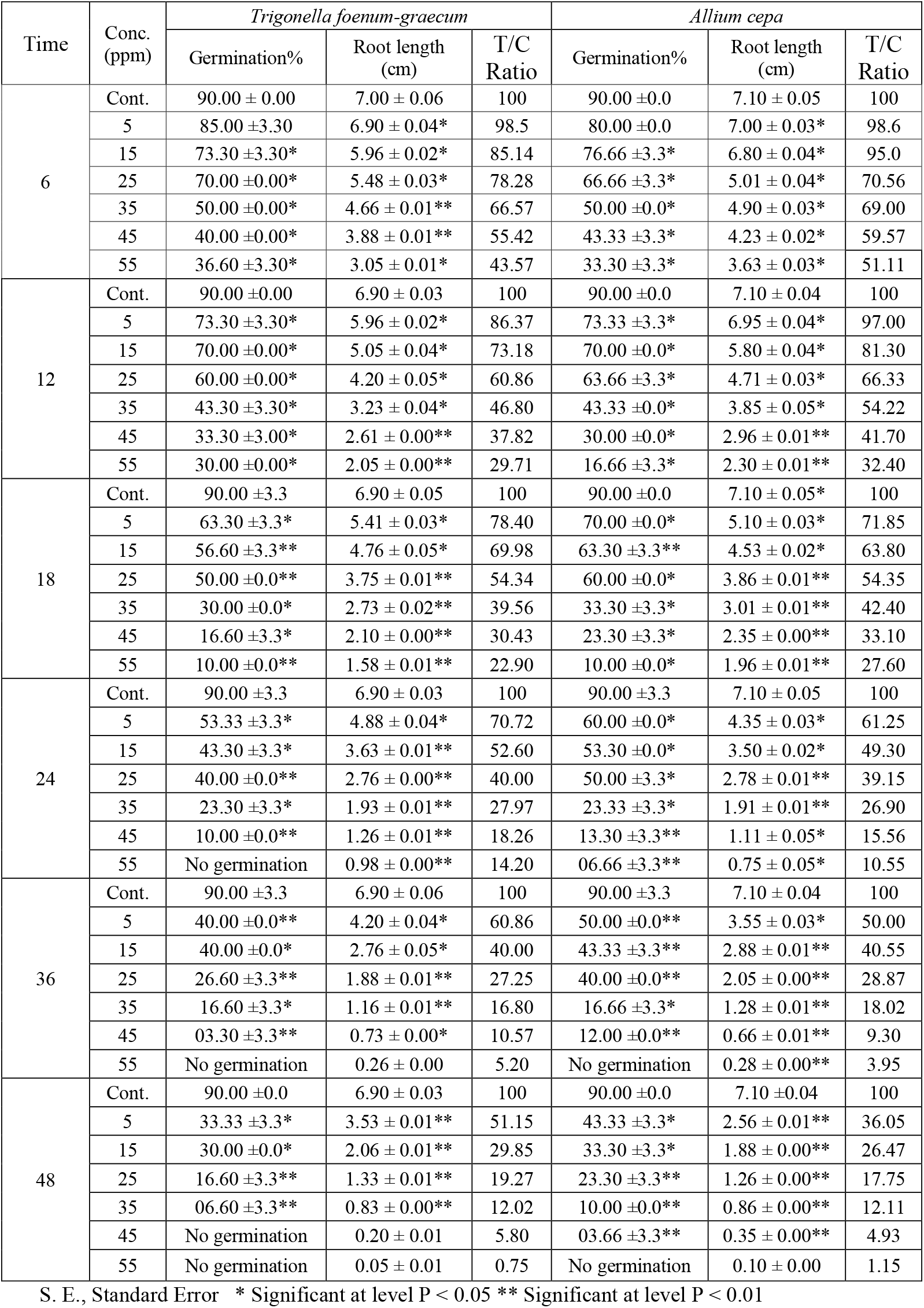
Seed germination percentage (%), radicle length (cm) and T/C ratio for *Trigonella foenum-graecum and Allium cepa* after treatments with different concentrations of maleic hydrazide.

| Time | Conc. (ppm) | *Trigonella foenum-graecum* |  |  | *Allium cepa* |  |  |
| --- | --- | --- | --- | --- | --- | --- | --- |
|  |  | Germination% | Root length (cm) | T/C Ratio | Germination% | Root length (cm) | T/C Ratio |
| 6 | Cont. | 90.00 ± 0.00 | 7.00 ± 0.06 | 100 | 90.00 ± 0.0 | 7.10 ± 0.05 | 100 |
|  | 5 | 85.00 ± 3.30 | 6.90 ± 0.04* | 98.5 | 80.00 ± 0.0 | 7.00 ± 0.03* | 98.6 |
|  | 15 | 73.30 ± 3.30* | 5.96 ± 0.02* | 85.14 | 76.66 ± 3.3* | 6.80 ± 0.04* | 95.0 |
|  | 25 | 70.00 ± 0.00* | 5.48 ± 0.03* | 78.28 | 66.66 ± 3.3* | 5.01 ± 0.04* | 70.56 |
|  | 35 | 50.00 ± 0.00* | 4.66 ± 0.01** | 66.57 | 50.00 ± 0.0* | 4.90 ± 0.03* | 69.00 |
|  | 45 | 40.00 ± 0.00* | 3.88 ± 0.01** | 55.42 | 43.33 ± 3.3* | 4.23 ± 0.02* | 59.57 |
|  | 55 | 36.60 ± 3.30* | 3.05 ± 0.01* | 43.57 | 33.30 ± 3.3* | 3.63 ± 0.03* | 51.11 |
| 12 | Cont. | 90.00 ± 0.00 | 6.90 ± 0.03 | 100 | 90.00 ± 0.0 | 7.10 ± 0.04 | 100 |
|  | 5 | 73.30 ± 3.30* | 5.96 ± 0.02* | 86.37 | 73.33 ± 3.3* | 6.95 ± 0.04* | 97.00 |
|  | 15 | 70.00 ± 0.00* | 5.05 ± 0.04* | 73.18 | 70.00 ± 0.0* | 5.80 ± 0.04* | 81.30 |
|  | 25 | 60.00 ± 0.00* | 4.20 ± 0.05* | 60.86 | 63.66 ± 3.3* | 4.71 ± 0.03* | 66.33 |
|  | 35 | 43.30 ± 3.30* | 3.23 ± 0.04* | 46.80 | 43.33 ± 0.0* | 3.85 ± 0.05* | 54.22 |
|  | 45 | 33.30 ± 3.00* | 2.61 ± 0.00** | 37.82 | 30.00 ± 0.0* | 2.96 ± 0.01** | 41.70 |
|  | 55 | 30.00 ± 0.00* | 2.05 ± 0.00** | 29.71 | 16.66 ± 3.3* | 2.30 ± 0.01** | 32.40 |
| 18 | Cont. | 90.00 ± 3.3 | 6.90 ± 0.05 | 100 | 90.00 ± 0.0 | 7.10 ± 0.05* | 100 |
|  | 5 | 63.30 ± 3.3* | 5.41 ± 0.03* | 78.40 | 70.00 ± 0.0* | 5.10 ± 0.03* | 71.85 |
|  | 15 | 56.60 ± 3.3** | 4.76 ± 0.05* | 69.98 | 63.30 ± 3.3** | 4.53 ± 0.02* | 63.80 |
|  | 25 | 50.00 ± 0.0** | 3.75 ± 0.01** | 54.34 | 60.00 ± 0.0* | 3.86 ± 0.01** | 54.35 |
|  | 35 | 30.00 ± 0.0* | 2.73 ± 0.02** | 39.56 | 33.30 ± 3.3* | 3.01 ± 0.01** | 42.40 |
|  | 45 | 16.60 ± 3.3* | 2.10 ± 0.00** | 30.43 | 23.30 ± 3.3* | 2.35 ± 0.00** | 33.10 |
|  | 55 | 10.00 ± 0.0** | 1.58 ± 0.01** | 22.90 | 10.00 ± 0.0* | 1.96 ± 0.01** | 27.60 |
| 24 | Cont. | 90.00 ± 3.3 | 6.90 ± 0.03 | 100 | 90.00 ± 3.3 | 7.10 ± 0.05 | 100 |
|  | 5 | 53.33 ± 3.3* | 4.88 ± 0.04* | 70.72 | 60.00 ± 0.0* | 4.35 ± 0.03* | 61.25 |
|  | 15 | 43.30 ± 3.3* | 3.63 ± 0.01** | 52.60 | 53.30 ± 0.0* | 3.50 ± 0.02* | 49.30 |
|  | 25 | 40.00 ± 0.0** | 2.76 ± 0.00** | 40.00 | 50.00 ± 3.3* | 2.78 ± 0.01** | 39.15 |
|  | 35 | 23.30 ± 3.3* | 1.93 ± 0.01** | 27.97 | 23.33 ± 3.3* | 1.91 ± 0.01** | 26.90 |
|  | 45 | 10.00 ± 0.0** | 1.26 ± 0.01** | 18.26 | 13.30 ± 3.3** | 1.11 ± 0.05* | 15.56 |
|  | 55 | No germination | 0.98 ± 0.00** | 14.20 | 06.66 ± 3.3** | 0.75 ± 0.05* | 10.55 |
| 36 | Cont. | 90.00 ± 3.3 | 6.90 ± 0.06 | 100 | 90.00 ± 3.3 | 7.10 ± 0.04 | 100 |
|  | 5 | 40.00 ± 0.0** | 4.20 ± 0.04* | 60.86 | 50.00 ± 0.0** | 3.55 ± 0.03* | 50.00 |
|  | 15 | 40.00 ± 0.0* | 2.76 ± 0.05* | 40.00 | 43.33 ± 3.3** | 2.88 ± 0.01** | 40.55 |
|  | 25 | 26.60 ± 3.3** | 1.88 ± 0.01** | 27.25 | 40.00 ± 0.0** | 2.05 ± 0.00** | 28.87 |
|  | 35 | 16.60 ± 3.3* | 1.16 ± 0.01** | 16.80 | 16.66 ± 3.3* | 1.28 ± 0.01** | 18.02 |
|  | 45 | 03.30 ± 3.3** | 0.73 ± 0.00* | 10.57 | 12.00 ± 0.0** | 0.66 ± 0.01** | 9.30 |
|  | 55 | No germination | 0.26 ± 0.00 | 5.20 | No germination | 0.28 ± 0.00** | 3.95 |
| 48 | Cont. | 90.00 ± 0.0 | 6.90 ± 0.03 | 100 | 90.00 ± 0.0 | 7.10 ± 0.04 | 100 |
|  | 5 | 33.33 ± 3.3* | 3.53 ± 0.01** | 51.15 | 43.33 ± 3.3* | 2.56 ± 0.01** | 36.05 |
|  | 15 | 30.00 ± 0.0* | 2.06 ± 0.01** | 29.85 | 33.30 ± 3.3* | 1.88 ± 0.00** | 26.47 |
|  | 25 | 16.60 ± 3.3** | 1.33 ± 0.01** | 19.27 | 23.30 ± 3.3** | 1.26 ± 0.00** | 17.75 |
|  | 35 | 06.60 ± 3.3** | 0.83 ± 0.00** | 12.02 | 10.00 ± 0.0** | 0.86 ± 0.00** | 12.11 |
|  | 45 | No germination | 0.20 ± 0.01 | 5.80 | 03.66 ± 3.3** | 0.35 ± 0.00** | 4.93 |
|  | 55 | No germination | 0.05 ± 0.01 | 0.75 | No germination | 0.10 ± 0.00 | 1.15 |

S. E., Standard Error \* Significant at level P < 0.05 \*\* Significant at level P < 0.01

The effect of MH on mitotic activity or mitotic index (MI) is given in Tables 2. In general, the MI values reduction in the treated roots of *Trigonella foenum-graecum* and *Allium cepa* was a dose and time dependent. For each treatment time, the reduction of MI increased with increasing MH concentrations from 5 to 55 ppm. The mitotic index was remarkably reduced in roots treated with 55 ppm for 48 h to a minimum value of 0.l8±0.02% in *Trigonella foenum-graecum* and to a value of 0.11± 0.02% in *Allium cepa*. Also, the results show the frequency of different mitotic stages varied according to the MH concentration for each treatment period (Table 2). In *Trigonella foenum-graecum* after 6h treatment, ana-telophase and metaphase frequencies were found to be elevated over the control values, which resulted in a substantial reduction in the prophase frequencies with the applied concentration to values below that of the control values. The frequency of prophase in treated root tips were decreased with increasing the concentration of MH for all treatment times. In *Allium cepa*, MH causes an increase in metaphase and ana-telophase frequencies with each treatment as compared with the control values. There is no significant effect on the frequencies of different mitotic stages following the treatment for 6 h with all concentrations of MH, but a significant and highly significant effect (P ≤ 0.05 and P ≤ 0.01) was recorded at 24, 36 and 48h with all concentrations of MH.

**Table 2.**
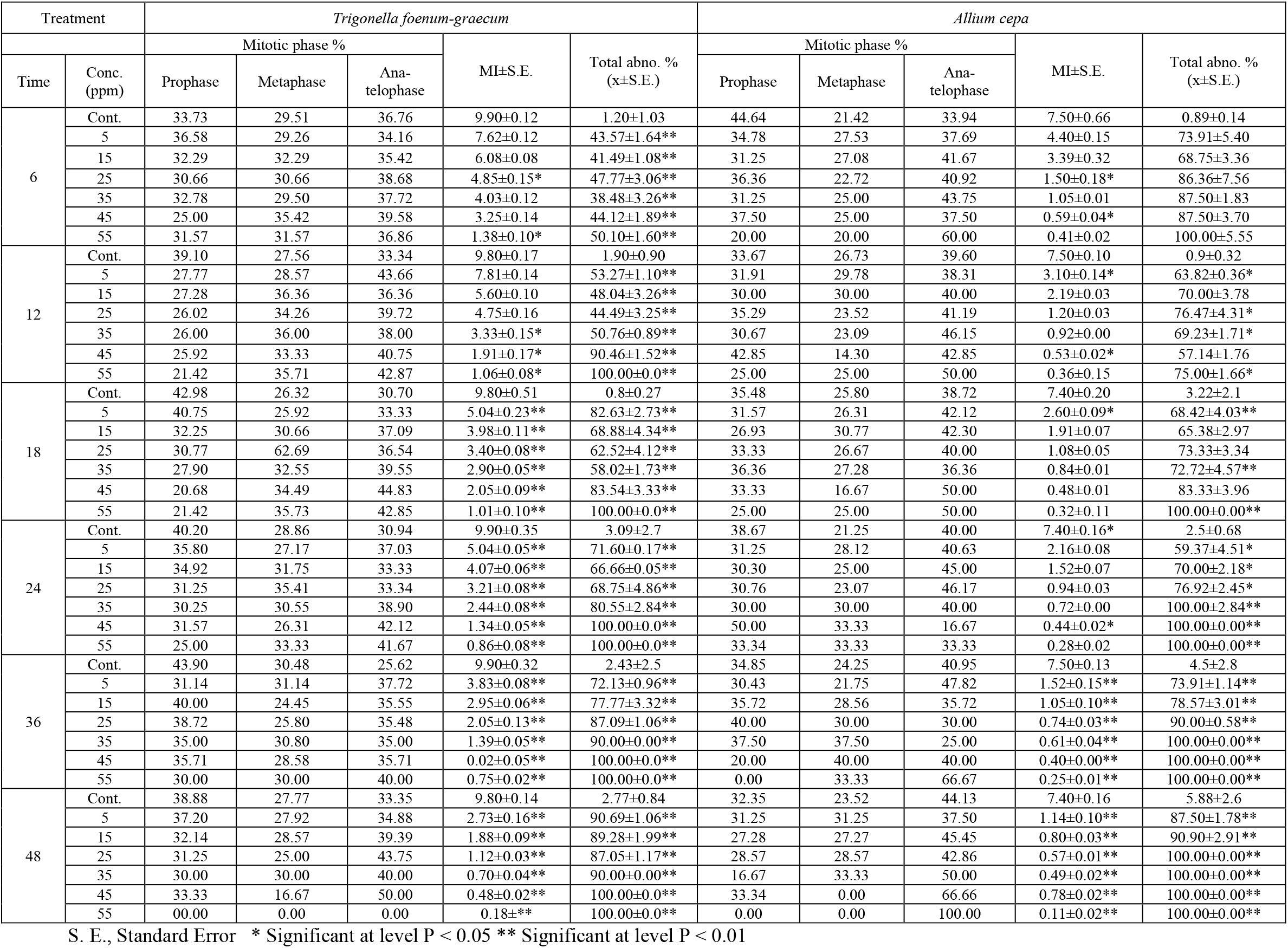
Mitotic index, percentage of mitotic phases and total mitotic abnormalities in *Trigonella foenum-graecum* and *Allium cepa* root-tip meristems after root-treatment with different concentrations of maleic hydrazide.

Result in Tables 3 and 4 shows that the total percentages of abnormalities in both test plants increased with increasing the concentration of MH and prolonged the treatment time and the induced different types of mitotic abnormalities in different stages of mitosis in root tips of both test plants following the treatments with MH. In *Trigonella foenum-graecum* the highest percentage of abnormalities was 100.0 ± 0.0% compared with the control value of 2.77±0.03 % which was recorded in roots treated with 55 ppm for 48 h. In roots treated of *Allium cepa* with 55 ppm for 48 all divided cells showed abnormalities (l00± 0.0%) compared with a control value of 5.88 ± 0.04%. Also, the results revealed a positive correlation was observed between the frequency of some types of abnormalities and both concentration of MH and treatment time. The ana-telophase stage appeared the most affected in both test plants with the highest percentages of abnormalities recorded for all treatment. The percentage of abnormalities induced at metaphase stage in *Allium cepa* was higher than that in *Trigonella foenum-graecum* for each treatment. In both plants, MH induced irregular prophase, vacuolated nuclei and sticky chromosomes, C-metaphase, bridges, disturbed metaphases and vagrants or laggards. In addition to the above-mentioned types, fragments, diagonal and forward chromosome(s) were recorded in low frequencies in treated root tip cells of both plants. Moreover, in *Allium cepa* only, star anaphase, micronuclei at interphase, multipolar and unequal sized nucleus were recorded with low frequencies. Some of these abnormalities are illustrated in Figures 1 and 2.

**Table 3.**
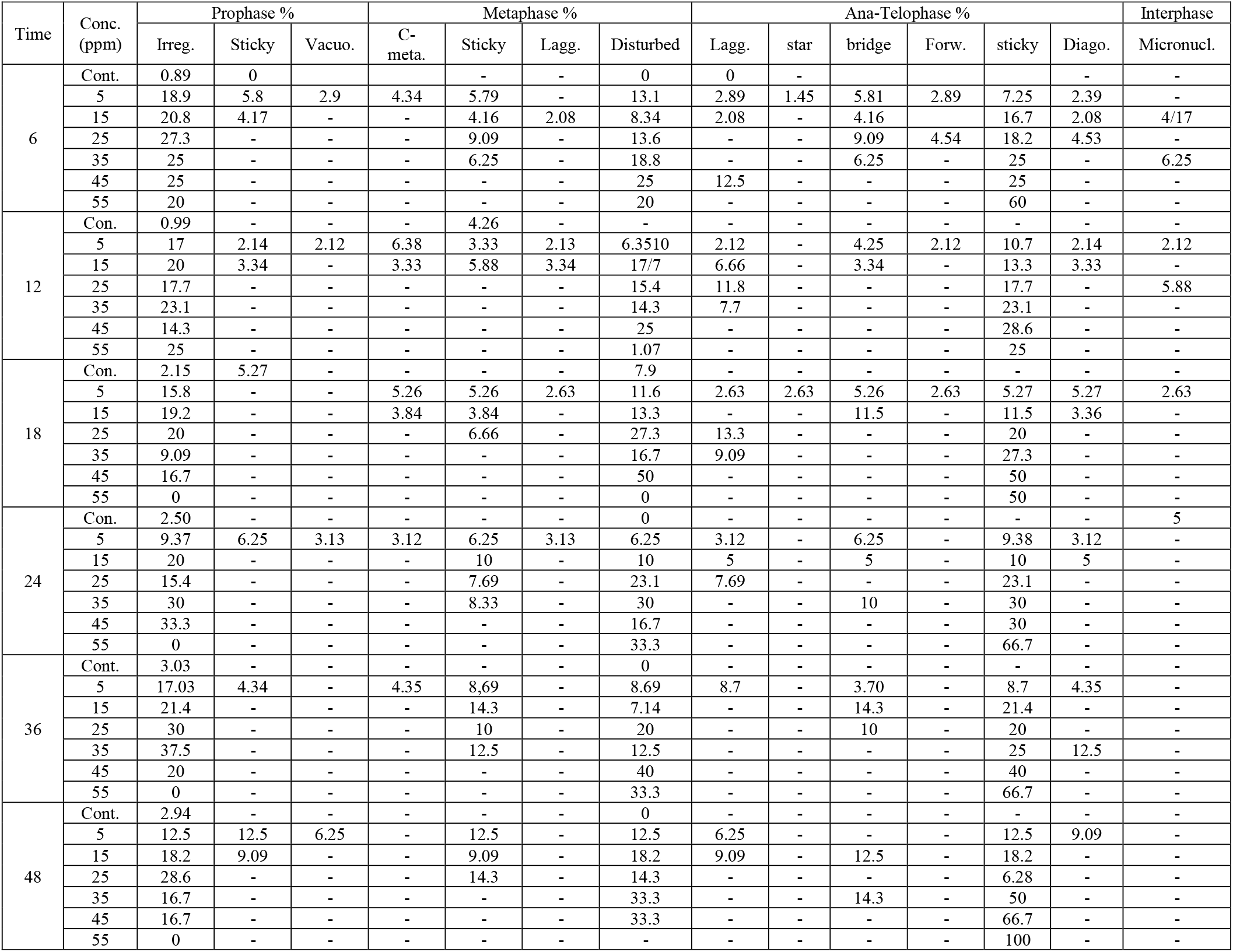
Types and percentage of mitotic abnormalities in different mitotic phases in *Allium cepa* root-tip meristems after treatments with different concentrations of maleic hydrazide for different times.

**Table 4.**
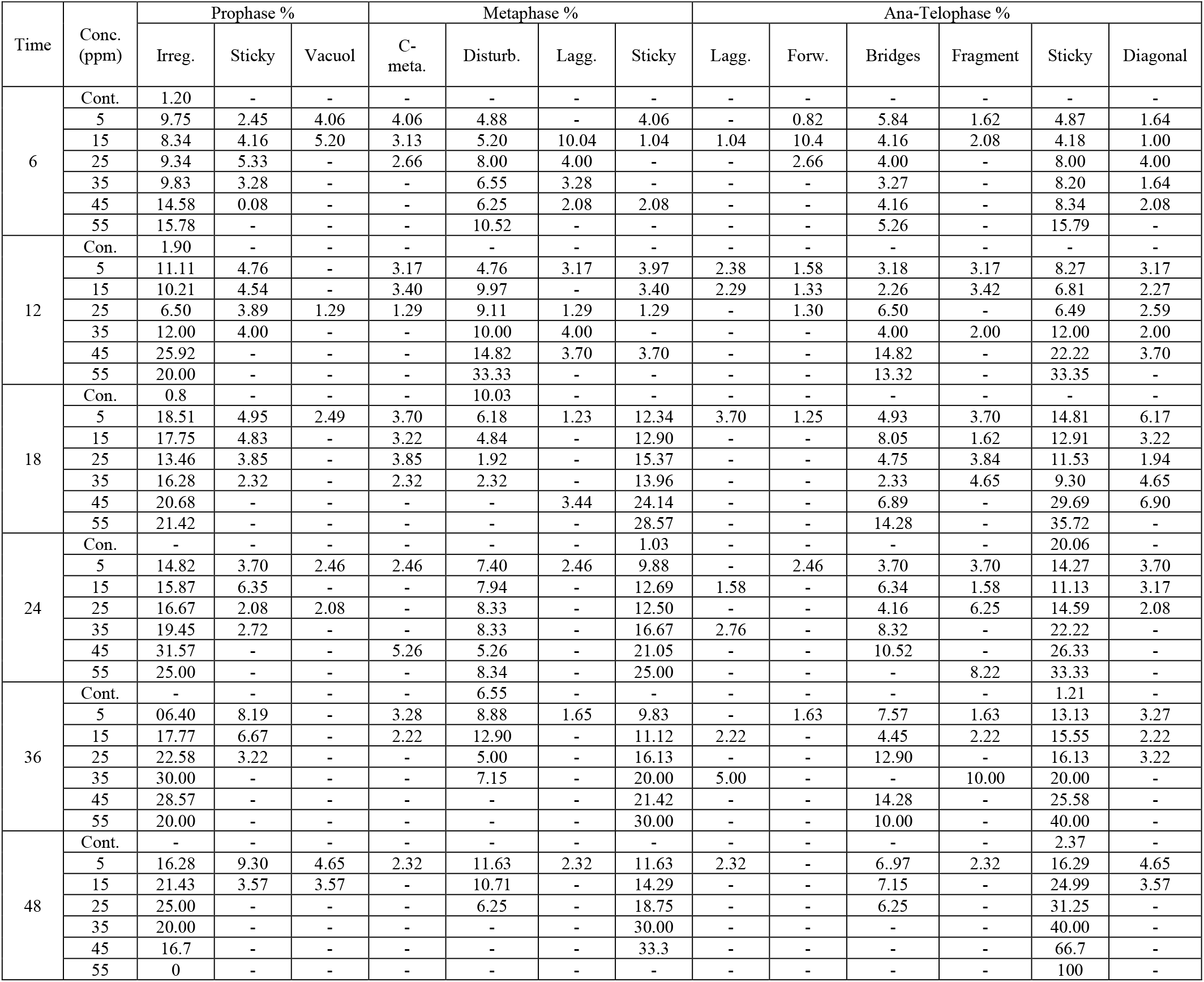
Types and percentage of mitotic abnormalities in different mitotic phases in *Trigonella foenum-graecum root*-tip meristems after treatments with different concentrations of maleic hydrazide for different times.

**Fig. 1.**
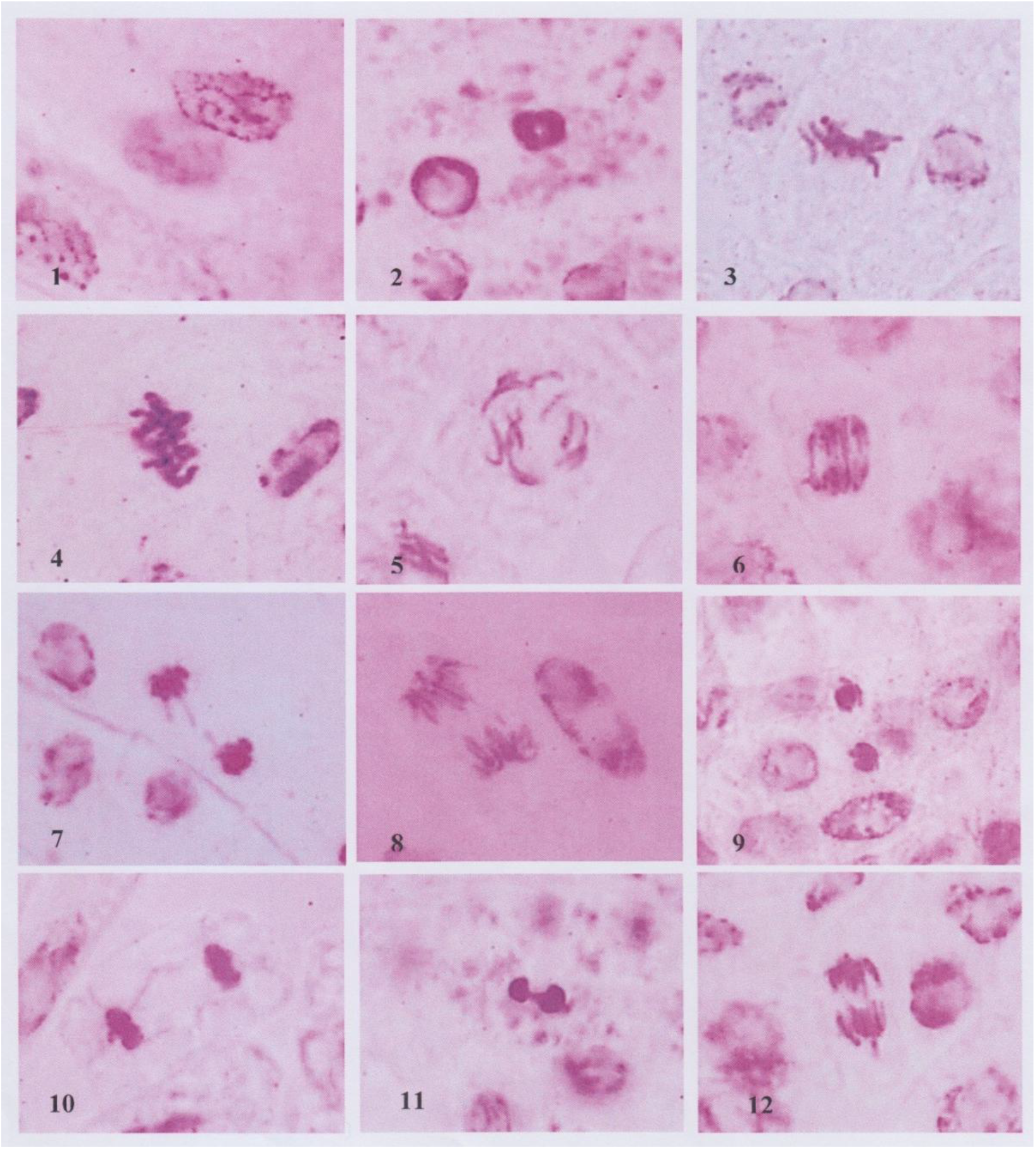
Some types of abnormalities induced in *Trigonella foanum-graecum* root tip cells by different treatments of MH. (1) Irregular prophase, (2) Sticky prophase, (3) Sticky metaphase with lagging chromosome, (4) Sticky metaphase, (5) C-metaphase, (6) Anaphase with multibridge, (7) Broken bridge at anaphase, (8) Anaphase with fragment, (9) Diagonal at telophase, (10) Sticky telophase with forward chromosome, (11) Sticky telophase with multibridge and (12) Anaphase with lagging and forward chromosome(s).

**Fig. 2.**
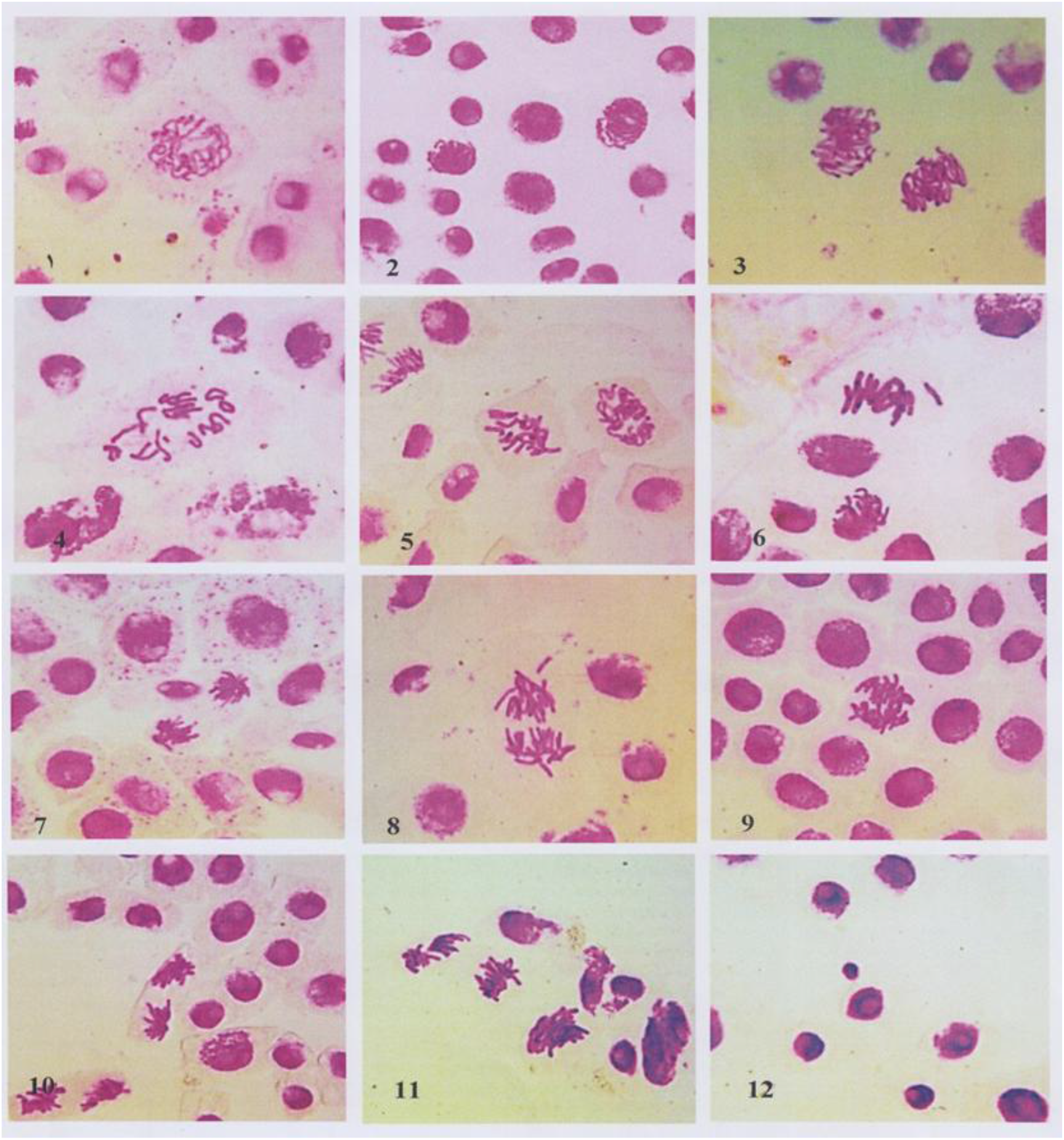
Some types of abnormalities induced in *Allium cepa* root tip cells by different treatments of MH.(1) Vacuolated nucleus at prophase, (2) Sticky prophase, (3) Irregular prophase, (4) C-metaphase, (5) Disturbed metaphase, (6) Sticky metaphase with lagging chromosome, (7) Star anaphase, (8) Anaphase with lagging and forward chromosome(s), (9) Multibridge at anaphase, (10) Diagonal at telophase, (11) Multipolar anaphase and (12) Micronucleus.

## Discussion

From the results both plants *Trigonella foenum-graecum* and *Allium cepa*, showed different response to maleic hydrazide effect with regard to seed germination percentage and radicle length obtained in this investigation. The effect of maleic hydrazide was dose dependent and more conspicuous at seedling stage affecting both root and shoot growth **(Kumar and Pal, 2004)**. The germination percentages in both plants were reduced by all applied concentrations of maleic hydrazide after prolonging the treatment time from 6 to 36h and were paralleled to its effects on root growth. Our results were in similar to that previously obtained by **Nickell (1953)** following treatment of *Rumex acetosa* seeds with maleic hydrazide and He concluded that the presence of MH caused the percentage of germination seeds of *Rumex acetosa* and root growth to be decreased and its effect was inhibitory rather than stimulatory and formative effects. So, this effect could be either by disturbance of the osmotic relationship which in turn affects some metabolic parameters during the early phases of germination as suggested by **Apelbaum and Burg (1972)**, or by germination injury, disturbance of physiological processes, suppression of cell division and induction of chromosomal aberration in dividing cells as mentioned by **Fiskesjo (1985), Padmavathi *et al*. (1992) and Sahi *et al*. (1998)**. Also, our results agreed with the finding of **(Agarwal and Ansari, 2001; De Marco *et al*., 2005; Jabee *et al*., 2008; Das *et al*., 2008 and Hasan *et al*., 2018)** following the treatment of different plants such as rice, corn, wheat, ragi, broad bean, fenugreek with maleic hydrazide and at 10^−2^ M concentration of MH root growth of *Helianthus annum* not occurred **(Kaymak, 2005)**. Moreover, **Venezian *et al*. (2017)** reported that MH demonstrated limited capacity to prevent seed germination *of Phelipanche aegyptiaca* where germination rate was not influenced by the lower MH application rates of 1.4-5.6 mM, but it was significantly reduced at 11.2 mM, and even more drastically at 22.4 mM (as compared to the untreated control) and all the germ tube of the germinated seeds at 11.2 and 22.4 mM were shorter than 3 mm.

In contrast to our results, **Grant and Harney (1960)** did not find any inhibitory effect on tomato seed germination at 10 mM MH, while *Albizia lebbeck* seeds responded in a positive germination rate to increase of MH concentration **(Tomar, 2008)**. In The failure of the treated seeds of both test plant in this study (fenugreek and onion) to germinate at high concentrations of maleic hydrazide may be a consequence of inhibited cell divisions and enlargements in the embryo and or an overall decrease in metabolic activity. The blockage of any one of the phases leading may completely inhibit the process of germination. So, according to the consideration of **Srivastava (1979)**, the reduction in germination percentage and root growth was due to weakening and disturbance of growth process resulting in early elimination of seedlings. Also, the inhibition of growth regulator and metabolic disturbance during germination may also be one of the reasons. It may be due to mutational changes at genetic or chromosomal level due to toxicity of MH as the reduction in germination corresponds with the increasing chromosomal aberrations **(Laskar and Khan, 2014)**. It may also be due to toxicity of MH followed by mutational changes at genic or chromosomal level because the reduction in germination corresponds with the increasing chromosomal aberrations **(Hasan *et al*., 2018)**.

Results in this study showed that the maleic hydrazide induced different mitotic changes in root tip cells of *Trigonella foenum-graecum and Allium cepa*. Such changes vary from (i) reduction of mitotic index of meristematic cells, (ii) changes in phase index and (iii) production of a large number of chromosomal aberrations. The inhibition of mitotic activity in both plants by MH is accompanied by reduction in seedling growth expressed as decreasing of radicle length or in other words the values of mitotic index reflected to the retardation of roots. This finding was similar to that reported by **(Mok, 1994; Osiecka and Janas, 1998; Das *et al*., 2008; Wismiowski *et al*., 2009 and Siddiqui *et al*., 2012)**. These changes appeared in varying degrees depending on duration of the treatment, where the maleic hydrazide caused gradually decreased in mitotic index, when compared with their respective controls. All the concentrations used induced highly significant effect (p < 0.01) at some treatments and significant effect at other treatments within 6, 12, 18 and 24 hours of treatment.

Generally, it has been observed that the extension of the treatment time and increasing the concentration of maleic hydrazide caused a retardation of cell division in the primary meristem of both plants when compared with their respective control values. The degree of mitotic inhibition is clearly dose dependent. Similar to our results, the reduction of mitotic activity seems to be a common effect of MH on mitosis in different plants as was mentioned by many investigators **(Cortes *et al*., 1985; Patil and Bhat, 1992; Edwin and Reddy, 1993; Rank and Nielsen, 1997; Osiecka and Janas, 1998; Mendhulkar, 2000; Kaymak, 2005; Sobita and Bhagirath, 2005; Mohanty *et al*., 2005; Juchimiuk, *et al*., 2007; Jabee, *et al*., 2008; Wismiowski *et al*., 2009; de Rainho *et al*., 2010; Pérez *et al*., 2011; Siddiqui *et al*., 2012 and Braszewska-Zalewska *et al*., 2014)**.

This mitotic inhibition could be attributed to one or more of these different causes such as: blocking of the mitotic cycle during interphase or at early prophase **(Juchimiuk *et al*., 2007)** or increasing the number of non-dividing cells **(Marcano *et al*., 2004)**, retarding the entrance of cells into the mitotic division by a marked prolongation of the cell cycle **(Dolezel *et al*., 1987)**, inhibition of spindle formation or cause spindle fibers defects and chromosome breakage during mitosis in root tips **(Rank *et al*., 2002 and Marcano *et al*., 2004)**, extending the duration of cell transitions from the S-phase of the cell cycle to mitosis **(Kwasniewska *et al*., 2018)**, blocking of cell cycle in the G_2_ phase, preventing the cell from entering mitosis, by the inhibition of DNA synthesis and/or protein synthesis or combining with protein to form chemical complex **(De Marco *et al*., 1992; Kaymak, 2005 and Debenest *et al*., 2008)**, inhibition of DNA replication **(Del Campo and Coletto, 1998)** or induction of DNA damage **(Alvarez-Moya *et al*., 2001; Menke *et al*., 2001; Juchimiuk *et al*., 2006; Ghosh *et al*., 2010 and Kwasniewska *et al*., 2018)**, inhibition the formation of various metabolic events necessary for mitosis **(Marcano *et al*., 2004 and Debenest *et al*. 2008)**. Later on, according to **Samuel *et al*. (2010)** a possible explanation for this is that with increasing concentration and consequently increasing toxic effect, there was an inhibitory effect on cell division. This might occur in pre-prophase, where cells are prevented from entering; prophase or there may be prophase arrest where cells enter into mitosis but are arrested during prophase resulting in a high frequency of prophase cells.

Moreover, in the light of its mode of action, maleic hydrazide presumably involved interference with sulphydryl groups which reduced the synthesis of those enzymes in which SH-groups were required, which can ultimately lead to metabolic disturbances in the cell **(Sanita di Toppi and Gabbrielli, 1999)**. The depression of the mitotic index observed in the present study could be brought about as a result of preventing cells from proceeding into prophase or from the depression of the mitotic phases following prophase stage. But the results obtained by **Kaymak (2005)** following the treatments of *Helianthus annuus* seeds with maleic hydrazide have shown that at 10^−2^ M concentration of MH root growth and mitotic division were not occurred and **Stroyev (1968)** reported that the high concentrations of MH completely suppressed cell division in barley root tip cells. Its effectiveness depended on the cell cycle stage treated. The result suggested that possible molecular mechanisms involved in the action of MH **(Cortes *et al*., 1985)**. To determine the mechanism through which the herbicide exerts its toxicity, ultra-structural electron microscopy was conducted. The results reveal nucleolar alterations, suggesting an inhibitory effect of biosynthetic activity **(Marcano *et al*., 2004)**.

The results showed that the percentage of each stage of mitosis which prophase, metaphase, ana-telophase in root tip cells of both plants was declined or elevated with all concentrations of maleic hydrazide used. Also, it was clear that the percentage of mitotic phases decreased or increased with increasing the period of the treatment from 6h to 36h. All treatments in both plants with maleic hydrazide altered the percentage of different mitotic phases in comparison to the control by increasing the percentages of prophase (which being the most common) and metaphase cells. The accumulation at prophase stage indicates that maleic hydrazide may be influenced on the chromosome spinalization or condensation and/or on the sequence of mitotic division and reducing the number of cells starting the division by blocking it at the end of prophase. This also could be probably due to the arrest in the cell cycle before metaphase to restore the integrity of DNA. It has been proposed that damaged DNA increase the time they stay at G_2_ and prophase, as a consequence of the working of checkpoints for unrepaired DNA, where post replication repair takes place **(Gonzalez-Fernandez and Lopez-Saez, 1985)**. With respect to the increased number of cells in metaphase in addition to prophase, this may suggest that may be mainly due to the interference with spindle formation, which causes an arrest of mitotic division at the end of the metaphase. This is clearly evident from the treatment with 50 and 55 ppm maleic hydrazide for 36h which stopped the mitotic division completely at metaphase stage in root tip cells in both plants. This may indicate that this chemical arrest mitosis at the end of the metaphase. These effects are characteristic of compounds which induce C-mitosis and are associated with the inhibitory action on formation of spindle by inhibition of ATPases which might be the cause of spindle disorganization **(Jabee *et al*., 2008)**.

The results in this investigation showed that the MH has a significant effect on the total percentages of the abnormalities induced in root tip cells in both plants and it was found to be positively correlated with the concentration and treatment time and was apparently different from that of the control. Chromosomal abnormalities constitute a significant portion of genetic damage produced by most mutagenic agents **(Kaymak, 2005)**. The total percentages of the abnormalities increased gradually with increase the MH concentration and as the period of treatment were prolonged as reported by **(Marcano *et al*., 2004 and Sabale and Mane, 2011 and Husain *et al*., 2013)**.

In general, the total percentages of mitotic aberrations induced by MH in *Allium cepa -* root tip cells were more than that induced in *Trigonella foenum-graecum*. The highest percentage of abnormalities induced by MH was recorded in divided cells at ana-telophase stages in both plants. All the applied concentrations of the plant growth regulator were capable to cause a wide range of percentages of chromosomal abnormalities in root tip cells of both tested plants and generally ranged from 57.14% to 100 % in *Allium cepa* and from 38.48% to 100% in *Trigonella foenum-graecum*. Similar observations of induction of chromosomal abnormalities have earlier been recorded by many workers tested the effect maleic hydrazide on different plants root tip cells such as **(Osiecka and Janas, 1998; Miadokové *et al*., 2001; Gichner, 2003; Marcano *et al*., 2004; Kaymak, 2005; Sobita and Bhagirath, 2005; Juchimiuk *et al*., 2006; Jabee *et al*., 2008; Debenest *et al*., 2008; Padmavthi *et al*., 2008; Ghosh *et al***.,**2010; Sabale and Mane, 2011 and Siddiqui *et al*., 2012)**.

Different types of mitotic abnormalities were observed in *Trigonella foenum-graecum* and *Allium cepa* after treatments with MH during the mitosis of root tip cells. The variation in the percentages of each of different types of chromosomal aberrations observed in this study was not dose dependent These types induced in both test plant may be grouped, according to **El-Ghamery *et al*. (2000)**, into: (a) mitotic abnormalities which indicates an effect on the mitotic apparatus such as C-mitosis, diagonal, forward chromosome, multipolar spindle formation, lagging and unequal distribution. Spindle disturbances may frequently be associated with defective movement of chromosomes. (b) chromosomal stickiness (at different stages), vacuolated nucleus, and irregular prophase and (c) chromosomal aberrations such as fragments, ana-telophase bridges and micronucleus at interphase, which were regarded to be due to an effect on the chemistry of chromosomes or chromosome structure involving chromosome breakage. However, these types could be physiological effects and clastogenic effects **(Nagpal and Grover, 1994)**. Also, **Saggoo *et al*. (2010)** classified abnormalities into: maximum clastogenic aberrations and non-clastogenic aberrations. In this concern, maleic hydrazide proved to be toxic as it produced bridges, laggards and micronuclei (clastogenic aberrations) in addition to non-clastogenic aberrations such as stickiness of chromosomes, vagrants, pycnosis and multi-polarity **(Sabale and Mane, 2000; De Marco *et al*., 2005 and Jabee *et al*., 2008)**. In this concern, **Sondhi *et al*. (2008)** reported that the wide spectrum of genotoxic effects induced by induced by maleic hydrazide (0.01%) in root tip cells of *Allium cepa* includes clastogenic (chromosomal bridges, chromosomal breaks) and physiological (stickiness, spindle inhibition, vagrant, multipolarity) aberrations.

Of these types of aberrations induced in this study, stickiness, as a most frequent type, was recorded in the different mitotic phases in root tip cells of both plants treated with MH. This type of abnormality was recorded by many investigators following the treatment of different plants by MH **(Edwin and Reddy, 1993; Sobita and Bhagirath, 2005; Jabee *et al*., 2008; Sondhi *et al*. (2008) and Siddiqui *et al*., 2012**,). Chromosome stickiness consisted of pro-metaphase sticky chromosomes and ana-telophases sticky bridges. The induction of this type reflected the toxicity of MH, which is most likely irreversible, and probably led to cell death **(Fiskesjo, 1985)**. Stickiness has been attributed to either the effect of MH on the denaturing activity of nuclear proteins or by physical adhesion involving mainly the proteinaceous matrix of chromatin material **(Patil and Bhat, 1992)**, which might also interfere with chromosome segregation (Gaulden, 1987) or binding with nucleic acids, mainly DNA, cause s serious changes in their physico-chemical properties **(El-Ghamery *et al*. 2000 & 2003)**, or chromatin condensation of the nucleus **(Klâsterskâ *et al*., 1976)**. Moreover, sticky chromosomes can cause loss of genetic material and subsequently, the cell-division précess occurs irregularly, with some chromosomes not adhering to the assembled chromosomal complex and being lost during the cell cycle.

The percentages of abnormal prophase in *Allium cepa* were higher than those in *Trigonella foenum-graecum* root tip cells. This type was a common type of abnormalities induced in treated root tip cells of both plants with MH and was recorded in a relatively high percentage in both plants. In this type, the chromatin threads were not typically arranged or the DNA in chromatin threads may be despiraled and/or depolarized. The induction of this type could be due to effect of MH on the process of individualization of chromatin threads to normal chromosome.

Another type of abnormalities at prophase was vacuolated nucleus which was in low and variable percentages in both plants and the percentage of this type was lower than the percentage of abnormal or irregular prophase. **El-Ghamery *et al*. (2003)** reported this type and concluded that chromatin lost its stain-ability or appeared as dense granulated, and nuclei become vacuolated. In this respect, **Khosla and Dnyansagar (1980)** observed chromatin granules scattered in the cytoplasm at telophase in *Cassia tora* root tip cells following the treatment with MH.

In this study, fragments were recorded in few cells in the root tips and were probably formed by acentric chromosome and as a result of inversion **(Agarawal and Ansari, 2001)**. Bridges lead to chromosome breakage and formation of acentric chromosome fragments which free-floating in the cytoplasm can be enveloped by membranes and produce micronuclei where a many cell was recorded with a micronucleus. Fragmentation might have arisen due to stickiness of chromosomes and consequent failure of separation of chromatids to poles **(El-Ghamery *et al*., 2003)**. According to **Leme and Marin-Morales (2009)** chromosome bridges and breaks in mitotic cells are indicators of a clastogenic action.

The induction of micronuclei indicates that some mitotic cells can enter into mitotic (M) phase with DNA damage. In this study, micronuclei, which probably, arose from the clastogenic action of MH as suggested by **(De Marco *et al*., 2005; Juchimiuk *et al*., 2007; Wiszniowski *et al*., 2009; de Rainho *et al*., 2010, Siddiqui *et al*., 2012 and Kwasniewska *et al*., 2018)**. Most micronuclei observed may have derived from a single chromosome that was lost from the whole set of chromosomes in the cell. This is related to the mechanism of action, namely MH can cause spindle fiber defects **(Mann *et al*. 1975; Jovtchev *et al*. 2001; Marcano *et al*., 2004 and Juchimiuk *et al*., 2007)**. Recently, The treatment of *Brachypodium distachyon* (Brachypodium) seeds with 3 mM or 4 mM MH for 3 hours caused a prominent clastogenic effect by induction of micronuclei in the cells at interphase and the frequency of the cells with micronuclei depended on the mutagen concentration **(Kus *et al*., 2017)**. In contrast to our results, the higher frequency of cells with micronuclei was observed after seedlings treatment of barley with 3mM MH **(Braszewska-Zalewska *et al*., 2014)** or in root meristematic cells in barley seeds induced by the same MH concentration, 3mM as shown by **(Juchimiuk *et al*. (2007)**.

The second frequent type of abnormalities was C-mitosis at (metaphase and anaphase stages) indicates either the action MH on the spindle fibers which involved in the processes of chromosome movement through regulation and control of de-polymerization and polymerization of the microtubules and preventing the continuation of the mitotic cycle **(Soh and Yong, 1993)** or the spindle apparatus was only impaired or it is one of the consequences of inactivation of spindle apparatus connected with the delay in the division of centromere. As a result of this disorder, the cell cycle is interrupted in metaphase and the chromosomes are seen scattered and condensed with very well-defined centromeres **(Fiskejo, 1985)**. These dose-independent changes were observed at all concentrations. The induction of some cells with disturbed metaphase chromosomes may suggest that MH caused partial disturbance in the spindle apparatus or partial suppression of spindle formation **(El-Ghamery *et al*., 2003)**. Consequently, a few cells with multipolar were also observed at ana-telophase referred to its effect by causing splitting of the spindle-fibers into more than two directions or partial suppression of spindle formation **(El-Ghamery *et al*., 2003)**.

In this study, the occurrence of bridges in *Trigonella foenum-graecum* than *Allium cepa* has been observed as similar to the results of **(Sobita and Bhagirath, 2005; Padmavathi *et al*., 2008; Jabee *et al*., 2008; Sondhi *et al*., 2008; Sabale and Mane, 2011, Siddiqui *et al*., 2012)**. The recorded bridges were not only anaphases with single bridge, but also, with two, three and multiple bridges. An increase of anaphases with more than one bridge was detected more frequently in higher MH concentrations. The decrease of anaphase with single bridge was probably due to the increase of the other anaphase bridge type. **Husain *et al*. (2013)** treated *Vicia faba* with MH and reported that bridges may have been produced due to sub chromatic exchanges, unequal exchange of dicentric chromosomes. In this study, Chromosome bridges may have occurred due to the chromosomal stickiness and subsequent failure of free anaphase separation.

Diagonal cell (depolarized ana-telophase) was another type of abnormalities observed in some treatments of MH in root tips of both plants. In this type, the two ana-telophase groups of chromosomes did not orient at the same axis of the cell. The formation of this type is indicative of the ability of substance to interfere with spindle apparatus mechanism.

Different types of abnormalities have been described as weak C-mitotic effects indicating risk of aneuploidy **(Fiskesjo, 1985)**, these types were forward chromosome(s) and (laggards or vagrant chromosomes). In the first type, the arms of chromosomes pointed out-word to the pole instead of centromere during the chromosome movement at ana-telophases. The other types are (laggards and vagrant chromosomes) are indicators of spindle poisoning **(Rank, 2003)**. Lagging chromosomes or laggards were observed in low percentage in both plants root tip cells treaded by MH in similar to that previously recorded by **(Jabee *et al*., 2008 and Sondhi *et al*., 2008)**. Lagging chromosomes are those that do not pinch out completely from their opposite daughter cell. The induction of this type might be also due to the delayed terminalization, stickiness of chromosomal ends or because of failure of chromosomes or acentric fragments to move to either pole **(Husain *et al*., 2013)**.

Vagrant chromosomes were observed in low percentage in root tip cells of *Allium cepa* treated by maleic hydrazide in similar to that previously recorded by **(Sondhi *et al*., 2008 and Siddiqui *et al*., 2012)**. These chromosomes were not organized to a specific stage of the mitotic division. This type of abnormality which is as a result of spindle irregularity or weakness of spindle formation, may be caused by unequal distribution of chromosomes with paired chromatids in which resulted from nondisjunction of chromatids in anaphase The induction of both types consequently leads to the separation of unequal number of chromosomes in the daughter nuclei and then formation of daughter cells with unequal sized or irregularly shaped nuclei at interphase **(El-Ghamery *et al*., 2003)**.

In conclusion, it was observed in the present study that the induction of cytogenetic effects using MH in the fenugreek and onion depends on the chemical doses and treatment time. In accordance with the previous data published for other species concerning the stronger cytogenetic effects induced by MH was in similar with our study. The amplitude of the responses of the studied plants to the action of MH as a plant growth regulator (growth retardant) was large enough to justify detailed studies concerning the genetic risks of its use.

